# Better together: a transboundary approach to brown bear monitoring in the Pyrenees

**DOI:** 10.1101/075663

**Authors:** Blaise Piédallu, Pierre-Yves Quenette, Ivan Afonso Jordana, Nicolas Bombillon, Adrienne Gastineau, Ramón Jato, Christian Miquel, Pablo Muñoz, Santiago Palazón, Jordi Solà de la Torre, Olivier Gimenez

## Abstract

Human administrative borders have no effect on wild animals, and the vast home ranges of large carnivores often cause them to live simultaneously on the territory of two or more countries or jurisdictions with different management policies. Here, we investigate the importance of transboundary population monitoring using as a case study the Pyrenean brown bear population (*Ursus arctos*) that lives in France, Andorra and Spain. Using capture-recapture models and the Pollock’s robust design, we estimated abundance and demographic parameters using data collected separately in France and Spain and a dataset gathered from joint monitoring on both sides of the border. As expected, the abundance estimates from French (from 11 bears in 2008 to 13 in 2014) or Spanish (from 4 bears in 2008 to 9 in 2014) data only were lower than abundance obtained from both sides of the border (from 11 in 2008 to 18 in 2014). The joint monitoring dataset also highlighted the importance of individual detection heterogeneity that, if ignored, would lead to underestimation. Our results reinforce the importance of transboundary cooperation when dealing with animal populations with territory spanning two or more administrative jurisdictions for collecting reliable scientific data and providing relevant abundance estimation to take sound management decisions.

## Introduction

Large carnivore populations in Europe are recovering from years of decrease after having been highly threatened or locally extinct, leading to larger population sizes and wider distributions (Chapron et al. 2014). These populations have large home ranges despite interspecific (Gittleman and Harvey 1982) or intraspecific (Mattisson et al. 2013; Nilsen et al. 2005) variations, and their recovery increases the likelihood of crossing the administrative border between two countries, or between two distinct jurisdictions within the same country. Therefore, the same large carnivore population may overlap several territories where the inhabitants have different or conflicting opinions on their monitoring or management (Bischof and Swenson 2012; Bull et al. 2009; Piédallu et al. 2016).

The conservation status of a population is commonly estimated based on its abundance and distribution, which in turn may help to assess its viability and to predict its range in the future. However, while transboundary large carnivore populations are extremely common, transboundary conservation policies are still rarely implemented (Gervasi et al. 2015; Linnell and Boitani 2012; Trouwborst 2010). Moreover, population monitoring is usually performed independently between the countries on one side and the other of the boundary. These countries in turn perform abundance estimation based solely on the data gathered on their own territory, leading to a discrepancy between management policy and ecological reality (Linnell and Boitani 2012; Trouwborst 2010). This approach might cause significant errors when trying to assess the abundance of a species, due to the possibility for a single animal to be seen across different territories (Bischof et al. 2016).

Here, we aimed at illustrating the value of collaborative monitoring at a transboundary scale, using brown bears (*Ursus arctos*) in the Pyrénées as a case study. Brown bears are widely distributed in the Northern hemisphere, being spread over North America (Waits et al. 1998), Europe (Swenson et al. 2011), and Asia (Hirata et al. 2013). The species as a whole is not endangered. However, some populations can be both small and isolated from others, leading to an important risk of local extinction (Linnell et al. 2008; Swenson et al. 2011). Among those endangered populations is the Pyrenean brown bear population, which resides in the mountains between Southwestern France and Northeastern Spain. Despite being reinforced by the reintroduction of individuals from Slovenia in 1996-97 (3 bears) and 2006 (5 bears) with another one scheduled for 2016 (1 bear), its viability is still considered to be low (Chapron et al. 2010)due to its small size, its fragmentation in different cores, and heavy inbreeding that might negatively impact its growth in a medium- or long-term future. Moreover, there are conflicting attitudes among local inhabitants over bear presence (Piédallu et al. 2016).

Combining data collected through camera trapping, systematic search-effort protocols and opportunistic observations in Spain and France, we estimated abundance and demographic parameters for the Pyrenean brown bear population using Pollock’s robust design capture-recapture models (Kendall et al. 1997). We reproduced the same analyses mimicking a lack of collaboration between the two countries and compared the results. Overall, we demonstrate the benefits of cooperation across the border in the monitoring and management of a large carnivore population.

## Material & Methods

### 1 Study area and bear population

This study was performed in the Pyrenees Mountains, on both sides of the border, which included Southwestern France, the North of the Spanish Autonomous communities of Catalonia, Aragon and Navarre, and the principality of Andorra (Figure 1). Bears that live in the Pyrenees descend from individuals that were translocated here from Slovenia in 1996-97 (2 females and 1 male) and 2006 (4 females and 1 male), although a hybrid between a Slovenian bear and a now-deceased Pyrenean female is still alive. The relocations were performed to save the population that was in critical danger of extinction in 1995, with only 5 individuals remaining, including a single adult female. The current population is split in two unconnected population cores(Camarra et al. 2015): one in the Western Pyrenees, over France, Aragon and Navarre, and one in Central-Eastern Pyrenees, over France, Catalonia, Aragon and Andorra.

**Figure 1:**
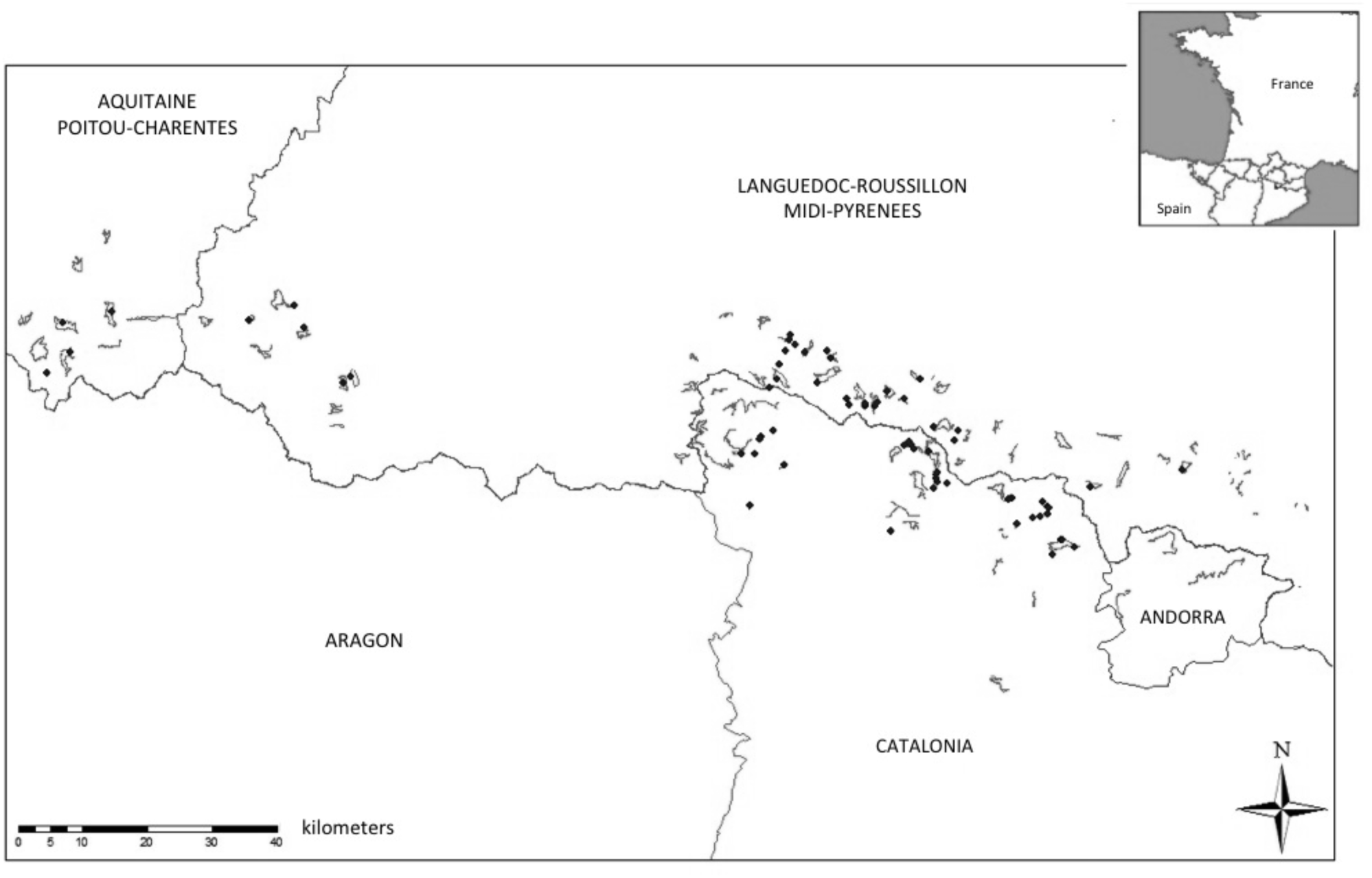
Map of bear monitoring in the French and Spanish Pyrenees. Bear monitoring in the French and Spanish Pyrenees.Thin lines: systematic itineraries. Dots: camera traps. The top-right mini-map indicates French counties and Spanish province – only the westernmost French county is part of the Aquitaine-Poitou-Charents region, all the others are part of the Languedoc-Roussillon-Midi-Pyrénées region. In France, data monitoring is handled by the Game and Wildlife Agency. In Spain, monitoring is handled separately between the three Autonomous communities: in Catalonia, the Fauna and Flora Service of the Department of Territory and Sustainability is in charge with a local delegation to the General Council in the Val d’Aran; in Aragon, it is the Environmental Services Department that oversees bear management, and in Navarre, it is the Public Society for Environmental Management of Navarre. In Andorra, monitoring is handled by the Natural Heritage Department.

### 2 Data collection and monitoring

Systematic monitoring was performed in France, Catalonia and Andorra from 2008 to 2014, consisting in fixed itineraries along which agents of governmental wildlife looked for bear tracks with help on the French side of the Pyrenees from volunteers of the Brown Bear Network, whose members include 135 professional members (from various agencies) and 228 amateur members (who are not affiliated to any agency). In Catalonia, two teams of 10 professional members handle systematic monitoring with help from Forestry Agents. Visits of these itineraries were repeated at regular time intervals between April and November, with at least a monthly visit (Figure 1). Due to the very low number of bears on their territories, there was no systematic monitoring in Aragon and Navarre. In all areas, data was also obtained through opportunistic means, which included all sightings or tracks found outside of systematic monitoring and validated by the local agencies, or damages caused on livestock (sheep in particular) or beehives. Since 2008, both countries have worked to standardize their monitoring protocols in the framework decided by the Transborder Monitoring Group for Bears in Pyrenees (GSTOP).

The most common bear tracks included hair samples (with 4-5 hair traps being scattered along the itineraries to improve the chances of getting a sample), scats, footprints, claw marks, photographs, films, attacks on livestock, sightings and opened ant-hills. Hair traps were added on trees along the itineraries to increase the chances to obtain hair samples. We discarded the tracks that did not allow the identification of an individual, mostly due to a lack of a good enough genetic sample. We also paid a particular attention to estimate the interval in which the bear might have left the track (see Supplementary materials). Genetic samples (hairs and faeces) were analyzed using a Polymerase Chain Reaction (PCR) multitube approach with four repeats for each sample to avoid the risk of misidentification due to the low DNA quantity of some hair samples (Miquel et al. 2006; Taberlet et al. 1997). 13 microsatellites markers and 1 marker for sex were targeted by the PCR in order to identify the bear individuals. The results were then aggregated between all sources.

We assigned to each individual its age class at first capture (Cub: up to 1 year old, Juvenile: up to 3 years old, Adult: 4 years old and more) and its gender. The population was assumed geographically closed, i.e. no emigration or immigration between this population and another one outside the Pyrenees.

### 3 Abundance estimation

We used the Pollock’s robust design (Kendall et al. 1997) to estimate abundance while accounting for imperfect detection of individuals and temporary emigration. The robust design approach uses repeated captures in a short timeframe, called secondary occasions, within a single time step (a so-called primary occasion). The population is considered to be closed (without demographic processes such as births, deaths, immigrations and emigrations) between two consecutive secondary occasions, hence allowing the robust estimation of abundance corrected for imperfect detection, and open (with births, deaths, emigrations and immigrations) between two consecutive primary occasions. In this study, we used years as primary occasions of capture (7 in total, from 2008 to 2014) and the months from May to September as secondary occasions (5 in total).We chose these secondary occasions because no births occur in this time interval, and we did not include the first months of the year because very young cubs have a high risk of dying.

We built three capture-recapture datasets: one for French data (including Andorra), one for Spanish data (Catalonia, Aragon, Navarre), and one for the Combined (France and Spain) data. We tested three possible types of temporary emigration: no emigration, a random emigration (where the probability of being a temporary emigrant only depends on the status of the individual at a given capture occasion) and a Markovian emigration (where the probability of being a temporary emigrant depends on the status of the individual at a given time step and at the previous time step).We tested 4 possible effects on survival: no effect, a sex effect, an age effect, and an additive effect of sex and age. Finally, 6 possible effects on detection were considered: no effect, a sex effect, a time effect (where the probability of detection changed between primary occasions – years, in our study), a mixture effect (with two distinct classes or individuals to account for detection heterogeneity, e.g. Cubaynes et al. (2010)), and the additive effects of sex and time and sex and mixture. In total, we tested 72 different models (Supplementary materials).

We used the Akaike Information Criterion corrected for small sample size (AICc) to perform model selection (Burnham and Anderson 2002). For each dataset (France, Spain, Combined), we kept the models with a AICc weight > 0.01. We also model-averaged parameter estimates to account for uncertainty in selecting one single ‘best’ model. To obtain 95% confidence intervals on abundance, we used a non-parametric bootstrap in which individual capture histories from each dataset were resampled with replacement a hundred times (Buckland et al. 1997).

We used the program MARK (White and Burnham 1999)called from software R (RCoreTeam 2013) with the ‘RMark’ package (Laake 2013).

## Results

### 1 Descriptive statistics

7517 bear tracks were gathered in the Pyrenees between 2008 and 2014. The number of tracks that were usable in a robust design framework for the French, Spanish and Combined dataset by both being accurate enough and collected between May and September (see Supplementary data) is shown in Table 1, and amounted for 15 to 17% of the total tracks.

**Table 1:** Number of bear tracks found yearly. Number of bear tracks per primary occasion of capture/Year. Two sets of secondary occasions are depicted: May-Sep (from May to September) and Apr-Nov (from April to November). The Total and Identified number of tracks in each dataset are shown in Table 1 of the Appendix section.

|  |  | 2008 | 2009 | 2010 | 2011 | 2012 | 2013 | 2014 | Total | %Total<br>Tracks | %ID'd Tracks |
| --- | --- | --- | --- | --- | --- | --- | --- | --- | --- | --- | --- |
| France | All year | 99 | 58 | 159 | 189 | 228 | 203 | 162 | 1098 | 22.80% | 74.34% |
|  | May-Sep | 80 | 52 | 123 | 130 | 157 | 161 | 121 | 824 | 17.11% | 55.79% |
|  | Apr-Nov | 94 | 56 | 154 | 178 | 208 | 199 | 147 | 1036 | 21.52% | 70.14% |
| Spain | All year | 37 | 54 | 90 | 145 | 101 | 48 | 93 | 568 | 21.02% | 63.18% |
|  | May-Sep | 24 | 41 | 65 | 124 | 66 | 29 | 77 | 426 | 15.77% | 47.39% |
|  | Apr-Nov | 36 | 51 | 80 | 142 | 101 | 48 | 91 | 549 | 20.32% | 61.07% |
| Total | All year | 136 | 112 | 249 | 334 | 329 | 251 | 255 | 1666 | 22.16% | 70.12% |
|  | May-Sep | 104 | 93 | 188 | 254 | 223 | 190 | 198 | 1250 | 16.63% | 52.61% |
|  | Apr-Nov | 130 | 107 | 234 | 320 | 309 | 247 | 238 | 1585 | 21.09% | 66.71% |

### 2 Model selection with the French dataset

The models that were selected for the French dataset mostly included a mixture effect on detection (Table 2A), whether alone or with an additive effect of sex. Survival was linked either to no effect, or to an effect of sex. However, no specific type of emigration (none, random or Markovian) was preferentially selected. The three best models (AICc weight > 0.1) included a sex effect along with individual heterogeneity on detection, and no effect on survival. No time effect on detection was found in the best models, and only the 12^th^ best had an age effect on survival.

**Table 2:** Model selection for each of the three datasets. List of the best models (Weight > 0.01) selected for each one of the three datasets ((a) France (b) Spain (c) Total). #: Number of the model. Survival, Detection, Emigration: effects tested on each parameter in the model. 0: No effect on the parameter. npar: number of parameters. Note: whenever model averaging was impossible due to the different nature of the models (with or without a mixture effect), we only selected the category with the highest combined weight.

| **(a) France** |  |  |  |  |  |  |  |
| --- | --- | --- | --- | --- | --- | --- | --- |
| # | Survival | Detection | Emigration | npar | AICc | Weight | Deviance |
| 6 | 0 | Sex + Mix | None | 12 | 649.3 | 0.27 | 636.1 |
| 30 | 0 | Sex + Mix | Markovian | 14 | 650.2 | 0.17 | 632.4 |
| 54 | 0 | Sex + Mix | Random | 13 | 651.1 | 0.11 | 635.6 |
| 12 | Sex | Sex + Mix | None | 13 | 651.2 | 0.10 | 635.7 |
| 28 | 0 | Mix | Markovian | 13 | 651.6 | 0.08 | 636.1 |
| 4 | 0 | Mix | None | 11 | 652.7 | 0.05 | 641.7 |
| 60 | Sex | Sex + Mix | Random | 14 | 653.1 | 0.04 | 635.3 |
| 34 | Sex | Mix | Markovien | 14 | 653.5 | 0.03 | 635.7 |
| 52 | 0 | Mix | Random | 12 | 653.8 | 0.03 | 640.6 |
| 10 | Sex | Mix | None | 12 | 654.5 | 0.02 | 641.2 |
| 26 | 0 | Sex | Markovian | 12 | 655.4 | 0.01 | 642.2 |
| 18 | Age | Sex + Mix | None | 19 | 655.5 | 0.01 | 625.9 |
| 58 | Sex | Mix | Random | 13 | 655.6 | 0.01 | 640.2 |
| 36 | Sex | Sex + Mix | Markovian | 15 | 655.8 | 0.01 | 635.7 |
| **(b) Spain** |  |  |  |  |  |  |  |
| # | Survival | Detection | Emigration | npar | AICc | Weight | Deviance |
| 49 | 0 | 0 | Random | 10 | 447.8 | 0.25 | 420.9 |
| 50 | 0 | Sex | Random | 11 | 448.1 | 0.21 | 418.9 |
| 55 | Sex | 0 | Random | 11 | 449.6 | 0.10 | 420.3 |
| 56 | Sex | Sex | Random | 12 | 450.0 | 0.08 | 418.4 |
| 25 | 0 | 0 | Markovian | 11 | 450.1 | 0.08 | 420.8 |
| 26 | 0 | Sex | Markovian | 12 | 450.5 | 0.07 | 418.8 |
| 52 | 0 | Mix | Random | 12 | 451.4 | 0.04 | 419.7 |
| 31 | Sex | 0 | Markovian | 12 | 451.9 | 0.03 | 420.3 |
| 54 | 0 | Sex + Mix | Random | 13 | 452.3 | 0.03 | 418.2 |
| 32 | Sex | Sex | Markovian | 13 | 452.4 | 0.03 | 418.3 |
| 58 | Sex | Mix | Random | 13 | 453.3 | 0.02 | 419.2 |
| 28 | 0 | Mix | Markovian | 13 | 453.7 | 0.01 | 419.7 |
| **(c) Combined** |  |  |  |  |  |  |  |
| # | Survival | Detection | Emigration | npar | AICc | Weight | Deviance |
| 12 | Sex | Sex + Mix | None | 13 | 725.0 | 0.21 | 724.5 |
| 6 | 0 | Sex + Mix | None | 12 | 725.2 | 0.19 | 726.9 |
| 58 | Sex | Mix | Random | 13 | 726.2 | 0.11 | 725.8 |
| 52 | 0 | Mix | Random | 12 | 726.4 | 0.10 | 728.1 |
| 18 | Age | Sex + Mix | None | 19 | 727.6 | 0.06 | 713.6 |
| 72 | Sex + Age | Sex + Mix | Random | 21 | 727.8 | 0.05 | 709.1 |
| 34 | Sex | Mix | Markovian | 14 | 728.4 | 0.04 | 725.8 |
| 28 | 0 | Mix | Markovian | 13 | 728.5 | 0.04 | 728.1 |
| 24 | Sex + Age | Sex + Mix | None | 20 | 728.6 | 0.03 | 712.3 |
| 10 | Sex | Mix | None | 12 | 728.9 | 0.03 | 730.7 |
| 64 | Age | Mix | Random | 19 | 729.2 | 0.02 | 715.3 |
| 4 | 0 | Mix | None | 11 | 729.4 | 0.02 | 733.3 |
| 36 | Sex | Sex + Mix | Markovian | 15 | 729.4 | 0.02 | 724.5 |
| 30 | 0 | Sex + Mix | Markovian | 14 | 729.6 | 0.02 | 726.9 |
| 70 | Sex + Age | Mix | Random | 20 | 730.2 | 0.02 | 713.9 |

### 3 Model selection with the Spanish dataset

All models selected for the Spanish dataset included emigration, whether it was random or Markovian (Table 2B), with the four best-ranked models having a random emigration. Models without emigration had a significantly lower weight. In contrast with the French dataset, the heterogeneity of detection (mixture effect) was seldom kept in the best models. Instead, most of the best models either had no effect or a sex effect for both survival and detection. The four best models (AICc weight >0.08) had a random emigration, and either no effect of a sex effect on both survival and detection.

### 4 Model selection with the Combined dataset

Like in the French dataset, all the best models for the Combined dataset had a mixture effect on detection, whether it was alone or with an additive effect of sex (Table 2C). No clear pattern emerged for emigration, although the models with Markovian emigration tended to have a lower weight. The results were similarly contrasted for the effects on survival, with sex being kept in 8 of the 15 models and age in 5 of 15.

### 5 Abundance estimates

In Spain, population increased to stabilize around 10 individuals, but showed a steep fall in 2013 before recovering in 2014 (Figure 2), which may indicate individuals that were temporarily unavailable. The French dataset pictured a very small increase between 2008 and 2014. The Combined dataset showed a trend similar to the Spanish dataset, with an increase between 2008 and 2014 and a temporary fall in 2013 mostly due to the lack of individuals detected on the Spanish side of the Pyrenees.

**Figure 2:**
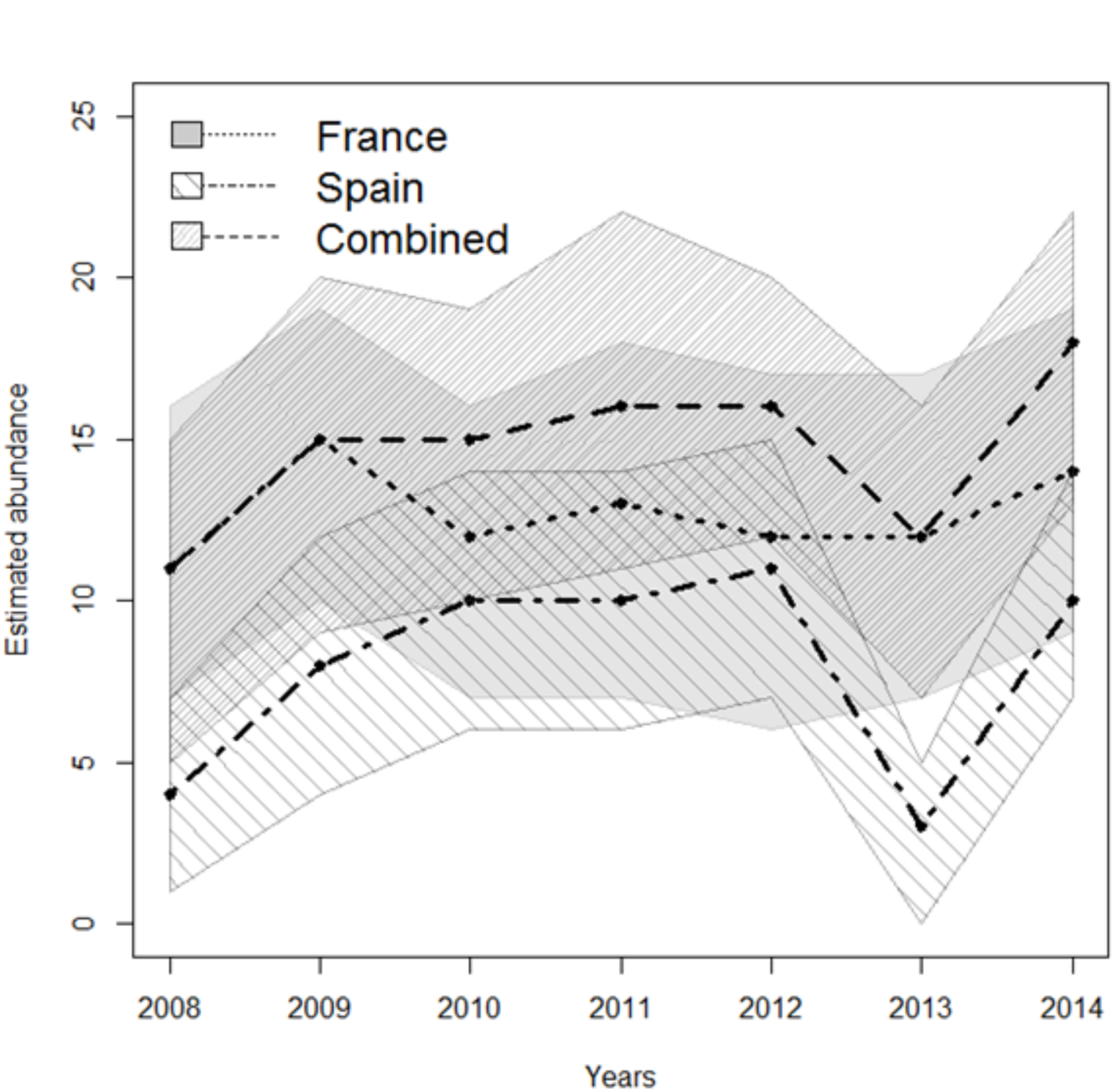
Population size estimation for each dataset. Population size for the Pyrenean brown bears between 2008 and 2014 estimated using the three different datasets (France only, Spain only, and Combined).

### 6 Demographic parameter estimates

The survival estimates were higher in France than in Spain (Table 3). There was a notable difference between male and female survival in the Combined dataset (F = 0.81±0.051, M = 0.933±0.041). In the France and Combined datasets, two classes of individuals were identified detection-wise, with one class being easily detected (~0.7-0.8) and one being harder to detect (~0.3-0.4). In both cases, the detection probability of females ended up being slightly inferior to the detection probability of males.

**Table 3:** Model-averaged parameter estimates. Model-averaged parameter estimates for each of the datasets. Bear Class represents the different individual classes identified by the mixture effect.

| Dataset | Bear Class | Survival |  | Detection |  | Emigration |  | Immigration |  |
| --- | --- | --- | --- | --- | --- | --- | --- | --- | --- |
| France | Female class 1 | 0.90 | $\pm 0.07$ | 0.84 | $\pm 0.13$ | 0.08 | $\pm 0.08$ | 0.50 | $\pm 0.46$ |
| | Female class 2 | | | 0.36 | $\pm 0.05$ | | | | |
| | Male class 1 | 0.92 | $\pm 0.06$ | 0.88 | $\pm 0.10$ | | | | |
| | Male class 2 | | | 0.45 | $\pm 0.06$ | | | | |
| Spain | Female | 0.83 | $\pm 0.06$ | 0.48 | $\pm 0.04$ | 0.27 | $\pm 0.09$ | 0.25 | $\pm 0.14$ |
| | Male | 0.85 | $\pm 0.07$ | 0.52 | $\pm 0.04$ | | | | |
| Total | Female class 1 | 0.88 | $\pm 0.05$ | 0.73 | $\pm 0.13$ | 0.07 | $\pm 0.05$ | 0.10 | $\pm 0.14$ |
| | Female class 2 | | | 0.32 | $\pm 0.05$ | | | | |
| | Male class 1 | 0.93 | $\pm 0.04$ | 0.81 | $\pm 0.05$ | | | | |
| | Male class 2 | | | 0.41 | $\pm 0.06$ | | | | |

In the Spain dataset, emigration was much higher than in the other datasets. This might be explained by the very few individuals found in 2013 that were probably treated by the models as temporary emigrating.

Moreover, all models selected for the Spain dataset included a random or Markovian emigration, while the France and Combined datasets also selected models with no emigration, which might have lowered the estimated values when averaged.

## Discussion

### 1 Abundance and heterogeneity in the detection process

The results for the French and Combined datasets highlighted two classes of individuals with significantly different detection. A previous study on wolves highlighted the importance of individual detection heterogeneity (IDH) when estimating abundance of large carnivore populations (Cubaynes et al. 2010). IDH in the Pyrenean brown bears might stem from intraspecific home range disparities (McLoughlin et al. 2000) making it more likely to find tracks of individuals who move a lot. The individual that was detected the most often, Pyros, is also the male with the highest reproductive success, and the three most commonly seen males (Pyros, Balou and Nere) were the three oldest living males in 2014. Another factor that might cause IDH is the efficiency of human agents when looking for bear tracks. Some Pyrenean bears display a stable spatial behavior over the years (Camarra et al. 2015), making their movements predictable in time. The personality differences displayed by bears (Fagen and Fagen 1996) might allow the agents to become better at finding tracks.

### 2 Sex and age effect on survival

A result that was consistent in all datasets was the lack of an age effect on survival. This is unexpected since young bears generally suffer from higher mortality rates (Bunnell and Tait 1985). However, because our analysis only included data ranging from May to September, we excluded the first four or five months of life for the bear cubs, during which their survival rate is at its lowest, meaning that bear cub deaths might have simply gone unnoticed due to the absence of the cubs from the dataset.

The presence of an effect of sex on survival, with higher mortality rates for females, seemed to contradict a previous study performed on grizzly bears where mortality was higher among males (McLellan et al. 1999). However, in this study, hunting explained the bias towards male mortality, since death from natural causes was more common among females. Because bear hunting is banned in the Pyrenees, the results between the two studies are actually consistent.

### 3 Estimates of the demographic parameters

Female bears survival in the Combined dataset was consistent with previous studies performed in other populations (De Barba et al. 2010; Garshelis et al. 2005; Wielgus and Bunnell 1994). Male bears survival in the same dataset was consistent with a previous study on a population without hunting pressure (De Barba et al. 2010). The emigration rate was significantly higher when using only the Spanish dataset compared to the French and Combined datasets, which could indicate that bears travel more from the Spanish to the French side of the Pyrenees. This important emigration rate on the Spanish side might also result from demographic stochasticity, stemming from the small size of the population and the very low number of individuals identified in Spain in 2013.

### 4 Robust design as a tool to estimate population size for cryptic species

Rare populations can either be defined by the low number of individuals, or by the fact that the population belongs to elusive, hard-to-observe species with large home ranges and low density (McDonald 2004). The Pyrenean brown bear population falls under both cases, and as such estimating its demographic parameters is made even more difficult. We confirmed however that robust design capture-recapture models allowed estimating several population parameters (Smith et al. 2013). In our case, the robust design study was combined with the use of multiple data sources (Boulanger et al. 2008; Gervasi et al. 2012) to obtain enough captures so we could build capture-recapture datasets with a sufficient number of observations to be exploited.

Abundance estimates were different from those calculated yearly by the GSTOP, which found a significant increase in the minimal population size between 2008 and 2014, going from 17 to 31 individuals (Camarra et al. 2015). This discrepancy can be explained by the method used by the agencies, which use both a Minimal Kept Population Size (EMR) and a Minimal Detected Population Size (EMD). Every year, the EMD can be used to correct the previous year’s EMR if some bears were not detected, due to the fact that the population is assumed geographically closed. In comparison, our robust design framework includes temporary emigration, which means that a bear that is not found during an entire year will not be included in the total population size. Moreover, to use the robust design framework, we excluded tracks that were difficult to date, and those that fell outside of the secondary occasions (May to September), which left some individuals identified by the GSTOP (Camarra et al. 2015) out of our database (see Supplementary materials). Agency estimates performed so far always included the individuals that were found dead in their yearly counts (Camarra et al. 2015), while a robust design model would only include them if the death occurred after the end of the primary occasion. Finally, the gender of a single individual born in 2012 was not determined, and as such that bear was excluded from our analyses. These factors put together account for the differences with the counts performed by the GSTOP.

### 5 Implications for bear abundance estimates

Even though different methods are used to estimate brown bear population size such as distance sampling and capture-recapture methods (Solberg et al. 2006), once the analyses are performed, most brown bear populations over the world are described by their total number of individuals, whether it is in Italy (Gervasi et al. 2008), in Sweden (Kindberg et al. 2011)or in the USA (Boulanger et al. 2008). Some other carnivore populations, however, are described with different indicators, such as the number of potential family groups (Andren et al. 2002) or the number of females with cubs of the year (Palomero et al. 2007). Considering the importance of the minimum population size for its viability (Shaffer 1981), we suggest that in complement to the minimal population size used by the GSTOP to describe the Pyrenean brown bears, it is important to develop an alternate counting method, especially since the heavy inbreeding (Camarra et al. 2015; Chapron et al. 2009) means that the genetic effective size (Palstra and Ruzzante 2008) of the Pyrenean brown bear population is lower than the “real” population size would let us to believe. Brown bear females in Europe (Swenson et al. 2007) usually start reproducing at the age of 4 or 5 with an interbirth interval of 2 years at least (Schwartz et al. 2003). Therefore, we suggest also describing the population by using the number of females with cubs of the year (Palomero et al. 2007) or the total number of 4+ years old females in the population, instead of limiting ourselves to the total number of bears. This would give a much more accurate insight of the viability of the population, which is an important tool to use when dealing with the conservation of endangered populations (Beissinger and Westphal 1998).

### 6 Importance of transboundary management

A transboundary management strategy helps to avoid significant errors in the estimates, most importantly the overestimation that occurs when a single individual is accounted for in one or more political jurisdiction due to its mobility(Bischof et al. 2016). The sum of yearly abundance in France and Spain was always larger than abundance from the Combined dataset, hence confirming this assumption. Even though administrative borders might coincide with a difference in local priorities (Moilanen and Arponen 2011), such as the Scandinavian lynx (*Lynx lynx*) population which is much more heavily hunted in Norway than it is in Sweden (Swenson and Andrén 2005),a transboundary approach allows for more biologically logical management (Linnell and Boitani 2012). Differing management policies between two neighboring jurisdictions can have consequences on a large carnivore population, e.g. by creating source/sink dynamics (Swenson and Andrén 2005)that will in turn affect demography (Robinson et al. 2008). Here, we showed that it was possible to build a transborder model despite differences in monitoring methods between the different jurisdictions.

## Conclusion

Perceived errors in abundance estimates might lead to skepticism towards the results provided by scientists, which can be followed by a negative evolution of a conservation conflict (Redpath et al. 2013). While accurate scientific data is not sufficient to solve the contention surrounding some large carnivore populations, it is a necessary step to advance towards mitigating conflicts by allowing the identification of human and ecological impacts (Redpath et al. 2013). It is especially important in the case of the Pyrenean brown bear, where the recent relocation of a male Slovenian bear in Catalonia could lead to an increase of the conflict, playing on the attitude differences between Pyrenean residents (Piédallu et al. 2016). In this context, reliable scientific data will be needed to avoid widening the gap between supporters and opponents of bear presence in the French-Spanish mountains. As large carnivore populations recover in Europe (Chapron et al. 2014), transboundary monitoring, however complex it might be (Kark et al. 2015),could become an essential first step towards harmonious and efficient transboundary management(Bischof et al. 2016; Kark et al. 2015), as it will be easier to implement than transboundary policies(Kark et al. 2015), and could facilitate the development of efficient solutions to ingrained conservation conflicts (Redpath et al. 2013).

